# Nuclear and mitochondrial genomic resources for the meltwater stonefly, *Lednia tumana* Ricker, 1952 (Plecoptera: Nemouridae)

**DOI:** 10.1101/360180

**Authors:** Scott Hotaling, Joanna L. Kelley, David W. Weisrock

**Affiliations:** School of Biological Sciences, Washington State University, Pullman, WA, USA; Department of Biology, University of Kentucky, Lexington, KY, USA

**Author notes:** Scott Hotaling, School of Biological Sciences, Washington State University, Pullman, WA, 99164, USA.

**Keywords:** Plecoptera, Nemouridae, *Lednia*, genomics, nuclear genome, USA

## Abstract

With more than 3,700 described species, stoneflies (Order Plecoptera) are an important component of global aquatic biodiversity. The meltwater stonefly *Lednia tumana* (Ricker, 1952; Family Nemouridae) is endemic to alpine streams of Glacier National Park and has been petitioned for listing under the U.S. Endangered Species Act (ESA) due to climate change-induced loss of alpine glaciers and snowfields. Here, we present *de novo* assemblies of the nuclear (~520 million base pairs [bp]) and mitochondrial (15,014-bp) genomes for *L. tumana*. The *L. tumana* nuclear genome is the most complete stonefly genome reported to date, with ~71% of genes present in complete form and more than 4,600 contigs longer than 10-kilobases (kb). The *L. tumana* mitochondrial genome is the second for the family Nemouridae and the first from North America. Together, both genomes represent important foundational resources, setting the stage for future efforts to understand the evolution of *L. tumana*, stoneflies, and aquatic insects worldwide.

## Introduction

Stoneflies are a diverse, globally distributed group of hemimetabolous insects that diverged from their closest relatives (e.g., Orthoptera, Dermaptera, Zoraptera) at least 300 million years ago in the Carboniferous Period (Béthoux, Cui, Kondratieff, Stark, and Ren 2011). With more than 3,700 described species, stoneflies account for a substantial portion of freshwater biodiversity (DeWalt, Kondratieff, and Sandberg 2015). The meltwater stonefly, *Lednia tumana* (Ricker, 1952; Plecoptera: Nemouridae), resides in alpine streams of Glacier National Park (GNP), USA, where it is iconic of habitat loss due to climate change in the region (Muhlfeld et al. 2011; Giersch, Hotaling, Kovach, Jones and Muhlfeld 2017). *Lednia tumana* is one of four extant species in the genus *Lednia* which all exhibit alpine, cold-water distributions in western North America (Baumann and Kondratieff 2010; Baumann and Call 2012). The majority of *L. tumana*’s habitat is supported by seasonal melting of permanent ice and snow, a habitat type that is under considerable threat of near-term loss as the global cryosphere recedes (Hotaling, Finn, Giersch, Weisrock, and Jacobsen 2017; Hotaling et al. 2019). The recent evolutionary history of *L. tumana* is closely tied to glacier dynamics with present-day genetic clusters arising in parallel with ice sheet recession at the end of the Pleistocene (~20,000 years ago, Hotaling et al. 2018). Genetic evidence has also highlighted a possible loss of mitochondrial genetic diversity for the species on even more recent, decadal timescales (Jordan et al. 2016). With such a narrow habitat niche in a small, mountainous region of the northern Rocky Mountains, *L. tumana* has been recommended for listing under the U.S. Endangered Species Act (US Fish & Wildlife Service 2016).

In this study, we present an assembly of the nuclear genome for *L. tumana*, the most complete nuclear genome for the order Plecoptera reported to date. This resource also represents one of only three high-coverage (> 50x) genomes for any EPT taxon (Ephemeroptera, *Ephemera danica*, Polechau et al. 2014; Plecoptera, this study; Trichoptera, *Stenopsyche tienmushanensis*, Luo, Tang, Frandsen, Stewart, and Zhou 2018), a globally important group of aquatic organisms commonly used for biological monitoring (e.g., Tronstad, Hotaling, and Bish 2016). We also present a nearly complete mitochondrial genome assembly (mitogenome) for *L. tumana*, the fourth for the stonefly family Nemouridae after two previous studies (Chen and Du 2017; Cao, Wang, Huang, and Li 2019).

## Materials and Methods

Genomic DNA was extracted using a Qiagen DNeasy Blood & Tissue Kit from a single *L. tumana* nymph collected in 2013 from Lunch Creek in GNP. Prior to extraction, both the head and as much of the digestive tract as possible were removed. A whole-genome shotgun sequencing library targeting a 250-bp fragment size was constructed and sequenced by the Florida State University Center for Genomics. The library was sequenced twice on 50% of an Illumina HiSeq2500 flow cell each time with paired-end, 150-bp chemistry, resulting in 242,208,840 total reads. The size of the *L. tumana* nuclear genome was estimated using a kmer-based approach in sga preQC (Simpson 2014). Read quality was assessed with fastQC (Andrews 2010) and low-quality reads were either trimmed or removed entirely using TrimGalore (Krueger 2015) with the flags: --stringency 3 --quality 20 --length 40. We assembled the nuclear genome using SPAdes v3.11.1 with default settings (Bankevich et al. 2012) and generated summary statistics with the Assemblathon2 perl script (assemblathon_stats.pl, Bradnam et al. 2013). The completeness of our nuclear assembly was assessed by calculating the number of conserved single copy orthologous genes [Benchmarking Universal Single-Copy Orthologs (BUSCOs)] in the assembly using BUSCO v3 and the 1,658 “insecta_ob9” set of reference genes (Simão, Waterhouse, Ioannidis, Kriventseva, and Zdobnov 2015). To compare the completeness of the *L. tumana* genome in the context of other stoneflies, we downloaded the two other publicly available stonefly genomes for *Isoperla grammatica* (Poda, 1761; Perlodidae) and *Amphinemura sulcicollis* (Stephens, 1836; Nemouridae) which are deposited under GenBank BioProject PRJNA315680 (Macdonald, Cunha, and Bruford 2016; Macdonald, Ormerod, and Bruford 2017). We performed the same BUSCO and Assemblathon2 analyses on the *I. grammatica* and *A. sulcicollis* genomes as we did for *L. tumana* above.

We assembled the *L. tumana* mitogenome with NOVOPlasty v2.6.7 (Dierckxsens, Mardulyn, and Smits 2016) using an 872-bp segment of the *L. tumana cytb* gene (GenBank KX212756.1) as the “seed” sequence. After assembly, the mitogenome was annotated through a combination of the MITOS web server with default settings (Bernt et al. 2013) and comparison to the *Nemoura nanikensis* mitogenome (Plecoptera: Nemouridae; Chen and Du 2017). However, in this initial assembly, the 16S gene was fragmented and the 12S gene was missing entirely. To mitigate this, we extracted sequences for both genes from the *N. nanikensis* mitogenome (Plecoptera: Nemouridae; Chen and Du 2017) and mapped our raw reads to these reference sequences for each gene with BWA-MEM v0.7.12-r1039 (Li 2013) using the default settings. Next, we used bcftools v1.9 (Li, Orti, Zhang, and Lu 2009) to collect summary information on the read mapping and genotype likelihoods (‘mpileup’ with default settings), called consensus nucleotides (‘call’ with -m flag), and output the consensus sequence (‘consensus’ with default settings). Finally, we used samtools v1.7 (Li et al. 2009) to calculate coverage depth per nucleotide for each sequence and masked consensus bases with no coverage (i.e., bases with no information from the *L. tumana* read mapping). We manually integrated our 16S and 12S sequences into the *L. tumana* mitogenome through comparison with the *N. nanikensis* mitogenome (Chen and Du 2017) and re-annotated the assembly with the MITOS web server (Bernt et al. 2013) as described above.

## Results and Discussion

The size of the *L. tumana* nuclear genome was estimated to be 536.7-megabases (Mb) from raw sequence data. Our final *L. tumana* genome assembly was 520.2-Mb with 50% of the assembly in contigs ≥ 4.69-kilobases (kb; Figure 1a, Table 1). The assembled genome size is in line with the only other publicly available stonefly genomes, *I. grammatica* (509.5 Mb) and *A. sulcicollis* (271.9 Mb). The *L. tumana* genome assembly also includes ~3,800 more contigs > 10 kb than the *A. sulcicollis* genome and ~4,600 more than the *I. grammatica* assembly (Figure 1a, Table 1). All associated data for the resources detailed in this study, including both raw reads and assemblies, are available as part of GenBank BioProject PRJNA472568 (mitogenome: MH374046, nuclear genome: SAMN09295077, raw reads: SRP148706).

**Figure 1.**
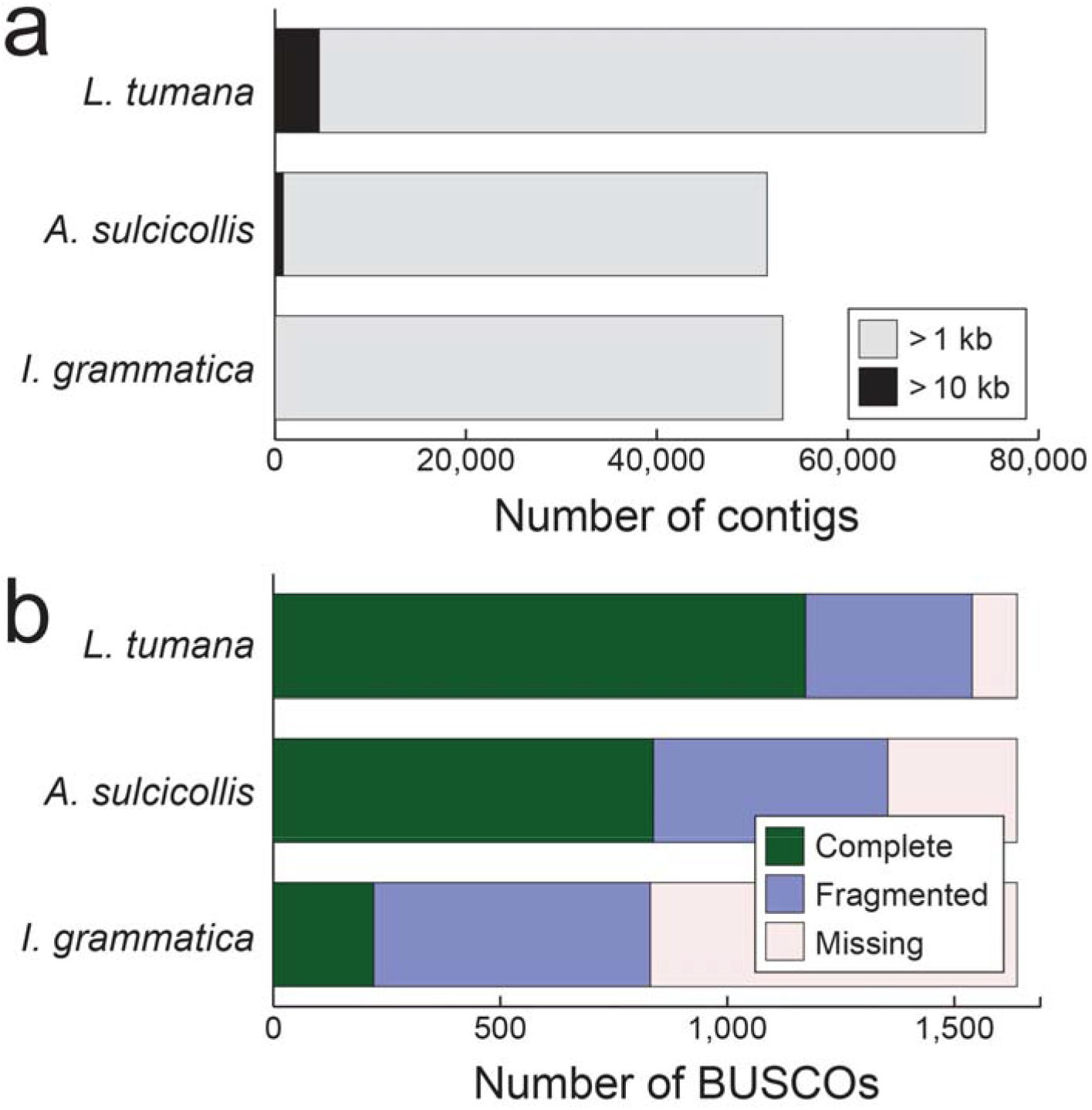
Comparisons of the gene content and contiguity of the *Lednia tumana* (Ricker, 1952; Plecoptera: Nemouridae) nuclear genome to the two other previously published stonefly genomes for *Amphinemura sulcicollis* (Stephens, 1836) and *Isoperla grammatica* (Poda, 1761; Macdonald et al. 2016; Macdonald et al. 2017). (a) The number of contigs > 1-kb and > 10-kb across the three stonefly genomes. For *I. grammatica*, the assembly contains just four contigs greater than 10-kb. (b) Presence of highly conserved, single-copy orthologous genes (BUSCOs) across the three stonefly genomes.

**Table 1.** Assembly statistics for the nuclear genome of *Lednia tumana* (Ricker, 1952; Plecoptera: Nemouridae) and two other stonefly species, *Amphinemura sulcicollis* (Stephens, 1836) and *Isoperla grammatica* (Poda, 1761; Macdonald et al. 2016; Macdonald et al. 2017). BUSCOs: Single-copy, orthologous genes known to be highly conserved among insects. A total of 1,658 BUSCOs were searched.

|  | *L. tumana* | *A. sulcicollis* | *I. grammatica* |
| --- | --- | --- | --- |
| Estimated genome size | 536,700,000 | n/a | n/a |
| Assembly size | 520,200,814 | 271,924,966 | 509,522,935 |
| Coverage | ~60x | ~1.4x | ~0.7x |
| Contigs > 1-kb | 74,445 | 51,555 | 53,204 |
| Contigs > 10-kb | 4,608 | 849 | 4 |
| Contig N50 | 4.69-kb | 0.85-kb | 0.46-kb |
| %A/T | 58.4 | 58.2 | 59.9 |
| %G/C | 41.5 | 42.8 | 40.1 |
| %N | 0.1 | 0 | 0 |
| Complete BUSCOs | 1,172 (70.6%) | 837 (50.5%) | 221 (13.3%) |
| Fragmented BUSCOs | 368 (22.2%) | 519 (31.3%) | 610 (36.8%) |
| Missing BUSCOs | 118 (7.2%) | 302 (18.2%) | 827 (49.9%) |

The meltwater stonefly’s nuclear genome is similarly A/T-rich (58.4%) to the other stoneflies (58–60%; Table 1), ants (55–67%; Gadau et al. 2012), *Drosophila melanogaster* (58%), *Anopheles gambiae* (56%), and the honeybee, *Apis mellifera* (67%, The Honeybee Genome Sequencing Consortium 2006). However, the *L. tumana* genome is far more complete in terms of genic regions than both existing stonefly assemblies with 92.8% of BUSCO reference genes either complete (70.6%) or fragmented (22.2%) versus 80.8% for *A. sulcicollis* (50.5% complete, 31.3% fragmented) and just 50.1% for *I. grammatica* (13.3% complete, 36.8% fragmented; Figure 1b, Table 1).

The *L. tumana* mitogenome is nearly complete, covering 15,014-bp, including all 13 protein-coding genes, 21 tRNA genes, both rRNA genes, and is only missing the control region (Figure 2). The organization of the *L. tumana* mitogenome is similar to that of *N. nankinensis*,the only other mitogenome available for the family Nemouridae. In *N. nankinensis*, the control region is ~1-kb which indicates that the complete *L. tumana* mitogenome is likely around 16-kb. This predicted size would fall in line with mitogenome sizes reported for other stoneflies (Chen and Du 2017). Our inability to resolve the control region is also unsurprising. In *N. nankinensis*, the control region contains a large ~1-kb repeat region which is inherently difficult to resolve without targeted long-range PCR re-sequencing or longer read high-throughput sequencing.

**Figure 2.**
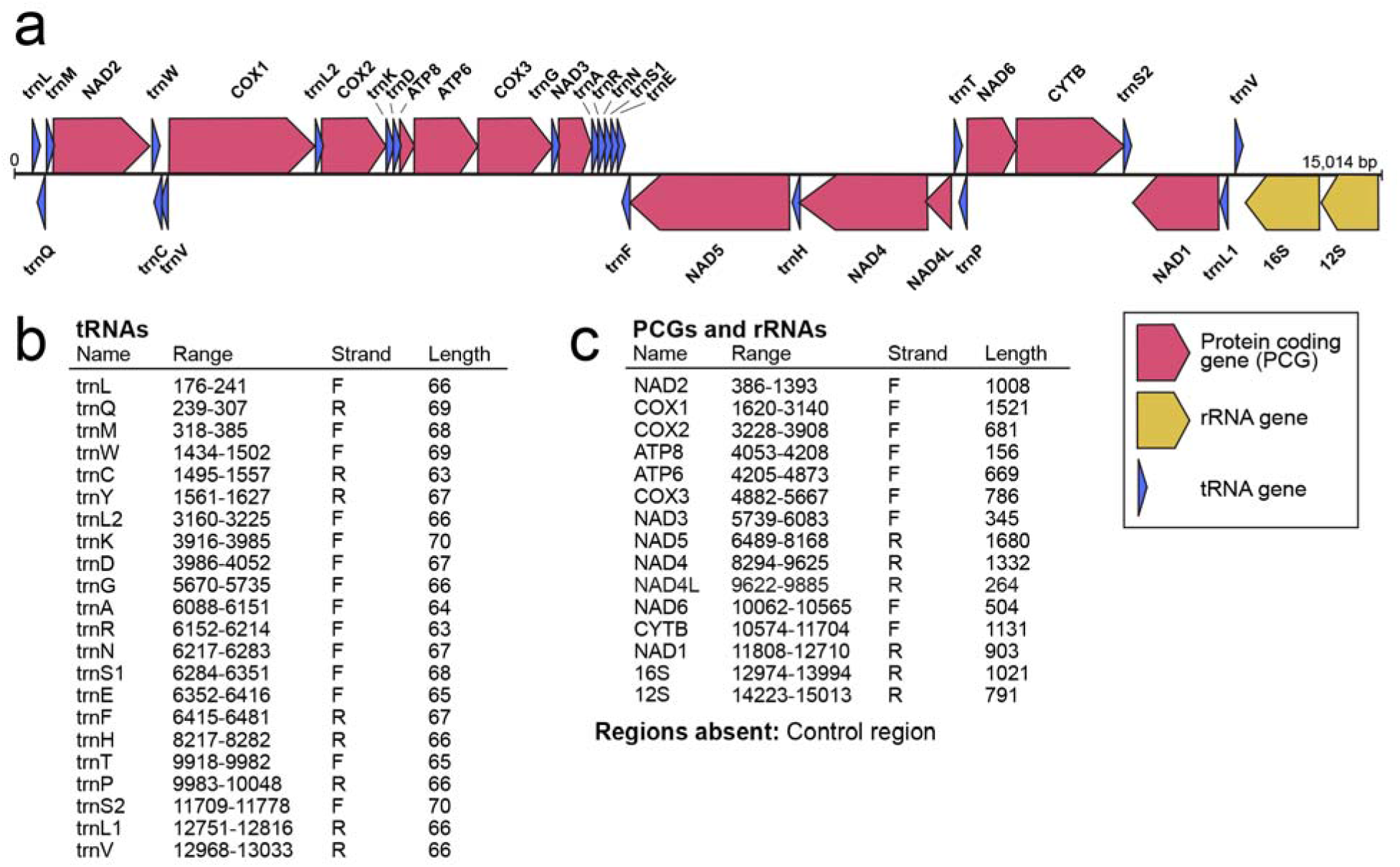
(a) The mitogenome of *Lednia tumana* (Ricker, 1952; Plecoptera: Nemouridae). (b) Locations of tRNAs. (c) Locations of protein coding (PCGs) and rRNA genes.

## Conclusion

The increasing availability of genome assemblies for a wide array of organisms is rapidly expanding the scope and comparative power of modern genome biology (Hotaling and Kelley 2019). With more than 4,600 contigs longer than 10-kb and ~70% of genes assembled in their complete form, the *L. tumana* nuclear genome provides new opportunity for exploring genome evolution within Plecoptera, a highly diverse, globally distributed insect order, or at higher levels of taxonomic organization (e.g., across all insects). Specifically, the *L. tumana* genome could be mined for genes for phylogenomic studies (e.g., Li et al. 2007; Borowiec, Lee, Chiu, and Plachetzki 2015) or more targeted, comparative assessments of specific genes or gene families across many taxa to clarify evolutionary shifts and/or copy number variation (e.g., Baalsrud et al. 2017).

Moreover, single-copy orthologous genes compared among species can provide a means for quantifying differences in evolutionary rates across diverse taxa (e.g., Honeybee Genome Sequencing Consortium 2006) and/or to identify rapidly evolving genes that underlie evolutionary transitions of interest. In the case of stoneflies and aquatic biodiversity in general, little is known of the evolutionary changes underlying the shift to an aquatic larval stage that is common among many orders (e.g., Plecoptera, Ephemeroptera, Trichoptera). With the addition of the *L. tumana* nuclear genome reported here to the recently published caddisfly *(S. tienmushanensis*, Order Trichoptera, Luo et al. 2018) and mayfly *(E. danica*, Order Ephemeroptera, Polechau et al. 2014) genomes, the stage is now set for broad, genome-scale investigations of how this major life history transition occurred across three globally distributed insect orders.

Future efforts to refine both assemblies, including the incorporation of longer reads (e.g., generated using Pacific Biosciences sequencing technology, Utturkar et al. 2014) will yield greater insight into the genome biology of *L. tumana*, stoneflies, and aquatic insects broadly. Still, the resources provided here, and particularly the most complete stonefly nuclear genome published thus far, represent an important step towards empowering modern stonefly research, a globally relevant group of aquatic insects that has been largely overlooked in the genomic age.

## Acknowledgements

We thank Alan Lemmon and Emily Lemmon for advice and assistance with sequencing. This research was supported by the University of Kentucky (UK) and National Science Foundation (DEB-0949532). Computational resources were provided by the UK Center for Computational Sciences and the Lipscomb High Performance Computing Cluster, as well as the Washington State University Center for Institutional Research Computing.

